# Preclinical Basis of the Efficacy and Pharmacodynamics of Finotonlimab, a Humanized Anti-PD-1 Monoclonal Antibody with Potent Implications for Clinical Benefit

**DOI:** 10.1101/2023.01.16.524197

**Authors:** Xiaoning Yang, Jing Li, Erhong Guo, Chunyun Sun, Xiao Zhang, Jilei Jia, Rui Wang, Juan Ma, Yaqi Dai, Mingjing Deng, Chulin Yu, Lingling Sun, Shuang Li, Liangzhi Xie

## Abstract

**Background:** The antibodies of programmed cell death protein 1 (PD-1) and its ligand (PD-L1) have dramatically changed the treatment landscapes for patients with cancer. Clinical uses of PD-1 antibodies have greatly improved the overall survival and durable responses in patients across selected tumor types.

**Methods:** We describe the preclinical characterization of Finotonlimab, a humanized anti-PD-1 antibody, by head to head comparison with Nivolumab or Pembrolizumab. Herein, we characterized the in vitro and in vivo efficacy, PK, PD and Fc mediated effector function of Finotonlimab. The single-agent anti-tumor activity of Finotonlimab was evaluated using humanized mouse models and a human PBMC reconstituted mouse model. Furthermore, in cynomolgus monkeys, comparative PK measurements confirmed better PK profiles of Finotonlimab than that of Pembrolizumab and Nivolumab.

**Results:** Our data showed Finotonlimab bind to human PD-1 with significantly high affinity and effectively inhibited its interaction with its ligands, PD-L1 and PD-L2, and thus could effectively stimulate the human T cell functions *in vitro* and exhibited significant antitumor efficacy *in vivo*. In addition, Finotonlimab showed minimal impact on Fc receptor dependent effector cell activation, which may contribute to the killing of PD-1^+^ T cells. In cynomolgus monkeys, Finotonlimab exhibited a non-linear pharmacokinetics (PK) profile in a dose-dependent manner, and approximately 90% of consistent receptor occupancy period was observed at 168 h after a single administration of 1 mg/kg. Following a 13-week successive administration of Finotonlimab, a pharmacodynamics study indicated a sustained mean receptor occupancy of ≥ 93% of PD-1 molecules on circulating T cells in cynomolgus monkeys up to 8 weeks even at 3 mg/kg.

**Conclusions:** Taken together, these preclinical data are encouraging and provide a basis for the efficacy and pharmacodynamics of Finotonlimab in clinical trials.

## Introduction

Targeting programmed cell death protein-1 (PD-1) and its ligand (PD-L1) has been an important immunotherapy strategy and has drastically changed the therapeutic landscape for patients with cancer. Anti-PD-1 antibodies have resulted in increased median overall survival and progress free survival in patients across multiple cancer types in clinical use [1, 2].

PD-1 is an inhibitory immune modulatory receptor [3-5]. It is expressed on activated T, natural killer cell (NK), B lymphocytes [6], macrophages, dendritic cells (DCs) [7], and monocytes [8] as an immune suppressor for both adaptive and innate immune responses [2-3]. Engagement of PD-1 by its ligands, PD-L1 [9] or PD-L2 [10, 11] leads to the exhaustion of T cell function and immune tolerance in the tumor microenvironment. Therefore, blockade of PD-1 pathway was considered a breakthrough to inhibit tumor immune escape and enhanced the T cell function to destroy the cancer cells, resulting in significant anti-tumor activity [12, 13].

Preclinical and clinical studies have demonstrated that antibodies targeting the PD-1/PD-L1 and blocking the receptor-ligand interactions could efficiently activate the T cell function and immune response [2]. However, the binding footprint between anti-PD-1 antibodies and PD-1 indicated different binding affinities and biological activities [14, 15]. The constant region of an antibody also plays an important role by exerting secondary pharmacodynamic effects through the binding to FcγRs or activation of complement cascade [16]. Therefore, the anti-PD-1 antibodies with different functional and pharmacokinetic characteristics have provided possibilities for different dosing requirements, safety concerns, and treatments for specific individuals and cancer types.

In this study, Finotonlimab (SCTI10A) was screened from a set of mAbs with high binding affinity from an antibody library by phage display and was humanized by grafting the heavy- and light-chain complementarity-determining regions onto the germline variable region frameworks of their nearest human species orthologs. Finotonlimab currently is being evaluated in several solid tumors in Phase III trials, including squamous-cell NSCLC (NCT04171284), HCC (NCT04560894), and HNSCC (NCT04146181), they all used as a monotherapy and combination with other drugs. Herein, we characterized the in *vitro* and in *vivo* pharmacology, Fc mediated effector function of Finotonlimab, and cynomolgus monkey PK and pharmacodynamics (PD). All these promising results provided a steppingstone for ongoing clinical trials.

## Results

### Finotonlimab has High Affinity and Binding Specificity to hPD-1

The association and dissociation diagrams of Finotonlimab to the human PD-1 protein were analyzed by bio-layer interferometry. It was found that Finotonlimab had a higher affinity to the hPD-1 protein with the K_D_ value of 6.48 × 10^−11^ M and a lower dissociation rate (1.95E-5 s^-1^) compared with Nivolumab (5.12E-5 s^-1^) (**Figure 1A-B, Table 1**). Finotonlimab had a relatively higher concentration-dependent binding ability to both hPD-1 protein and human PD-1-engineered Jurkat cells, compared with Nivolumab (**Figure 1C, 1D**). The EC_50_ values of Finotonlimab and Nivolumab to hPD-1 protein were 31.5 ng/mL and 179 ng/mL, respectively, and an increased EC_50_ fold of 5.68 of Finotonlimab was observed. In addition, Finotonlimab showed specific binding to PD-1 without binding to other CD28 homologous proteins, including CD28, CTLA-4, BTLA and PIGF (**Figure 1E**).

**Figure 1.**
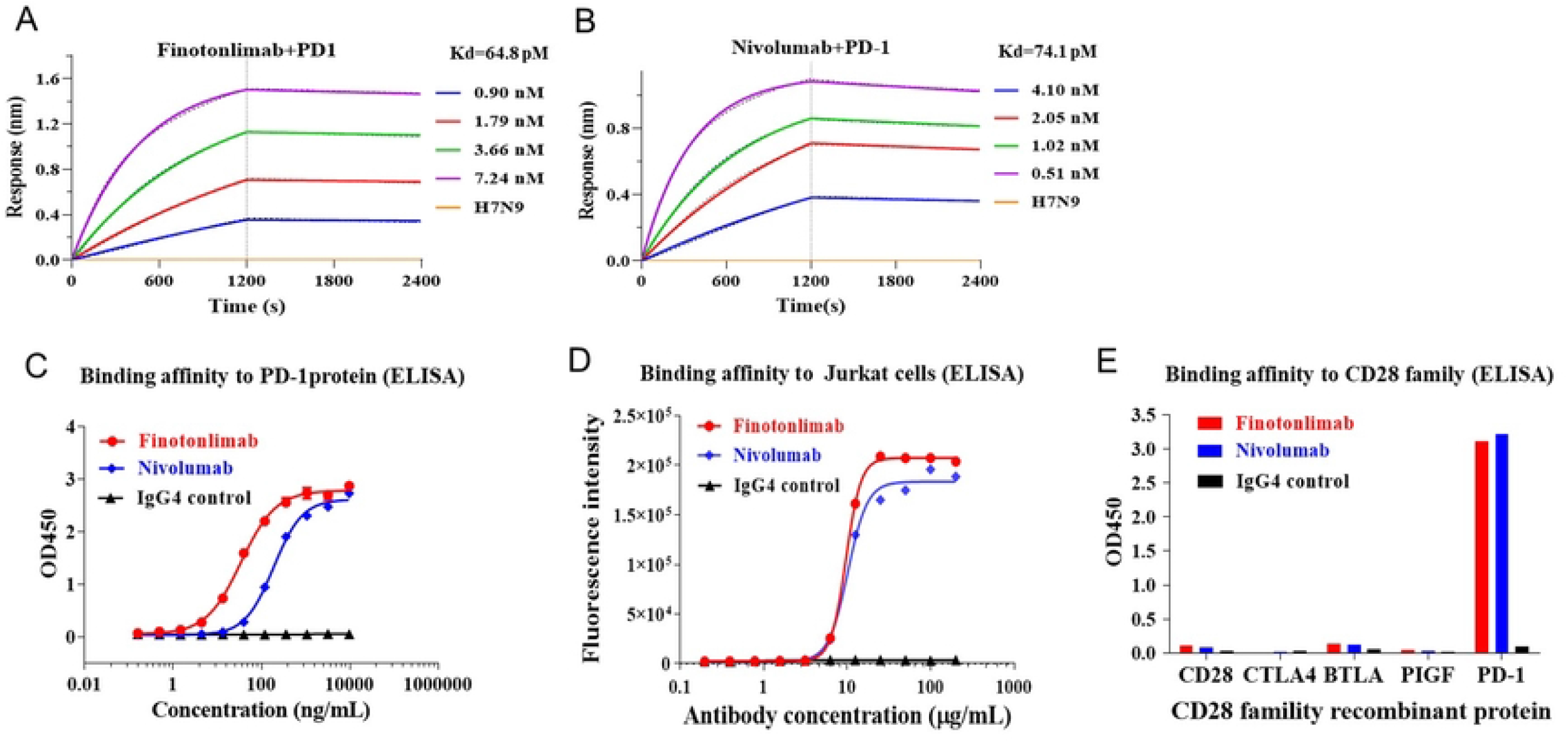
Affinity and binding specificity of Finotonlimab to hPD-1 by Octet and cell-based assays. (A). Binding affinity of Finotonlimab (A) and Nivolumab (B) to hPD-1 by bio-layerinterferometry (n=1). (C) Binding affinity of Finotonlimab and Nivolumab to hPD-1 protein by ELISA (n=3/group). (D) Binding affinity of Finotonlimab and Nivolumab to Jurkat cells. (E) Binding affinity of Finotonlimab and Nivolumab to CD28 homologous proteins, including CD28 and CTLA-4 (n=1).

**Table 1.**
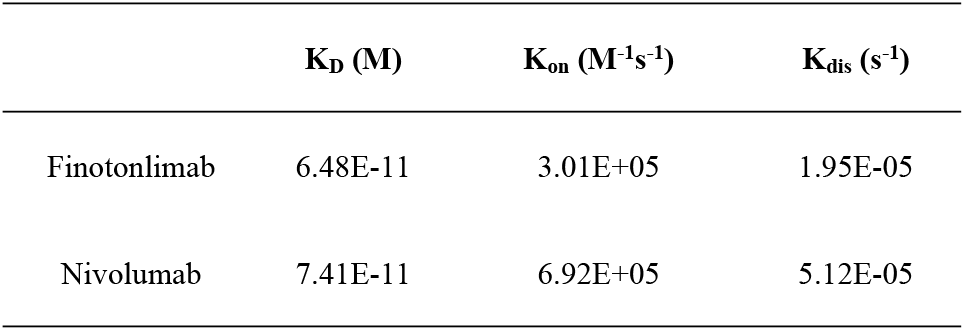
The binding affinity of Finotonlimab in comparison with Nivolumab.

| | $K_D$ (M) | $K_{on}$ (M<sup>-1</sup>s<sup>-1</sup>) | $K_{dis}$ (s<sup>-1</sup>) |
| --- | --- | --- | --- |
| Finotonlimab | 6.48E-11 | 3.01E+05 | 1.95E-05 |
| Nivolumab | 7.41E-11 | 6.92E+05 | 5.12E-05 |

### The Epitope of Finotonlimab Overlaps with the PD-L1/PD-L2 Binding Sites

The mutation experiment was performed to identify the epitope of Finotonlimab as well as the binding sites of PD-L1 (**Figure 2A**). The defined PD-1 mutants of N66, K78, K131, and E136/R139 had significantly reduced binding abilities (<40%) with PD-L1. These binding sites of PD-L1 indicate by our PD-1 mutants study were consistent with that analyzed by the PD-1 and PD-L1 complex structure (PDB ID 4ZQK) (**Figure 2B**). The binding sites of PD-L2 on PD-1 analyzed by the PD-1 and PD-L2 complex structure (PDB ID 6UMT) showed a similar pattern as that of PD-L1 (**Figure 2C**).

**Figure 2.**
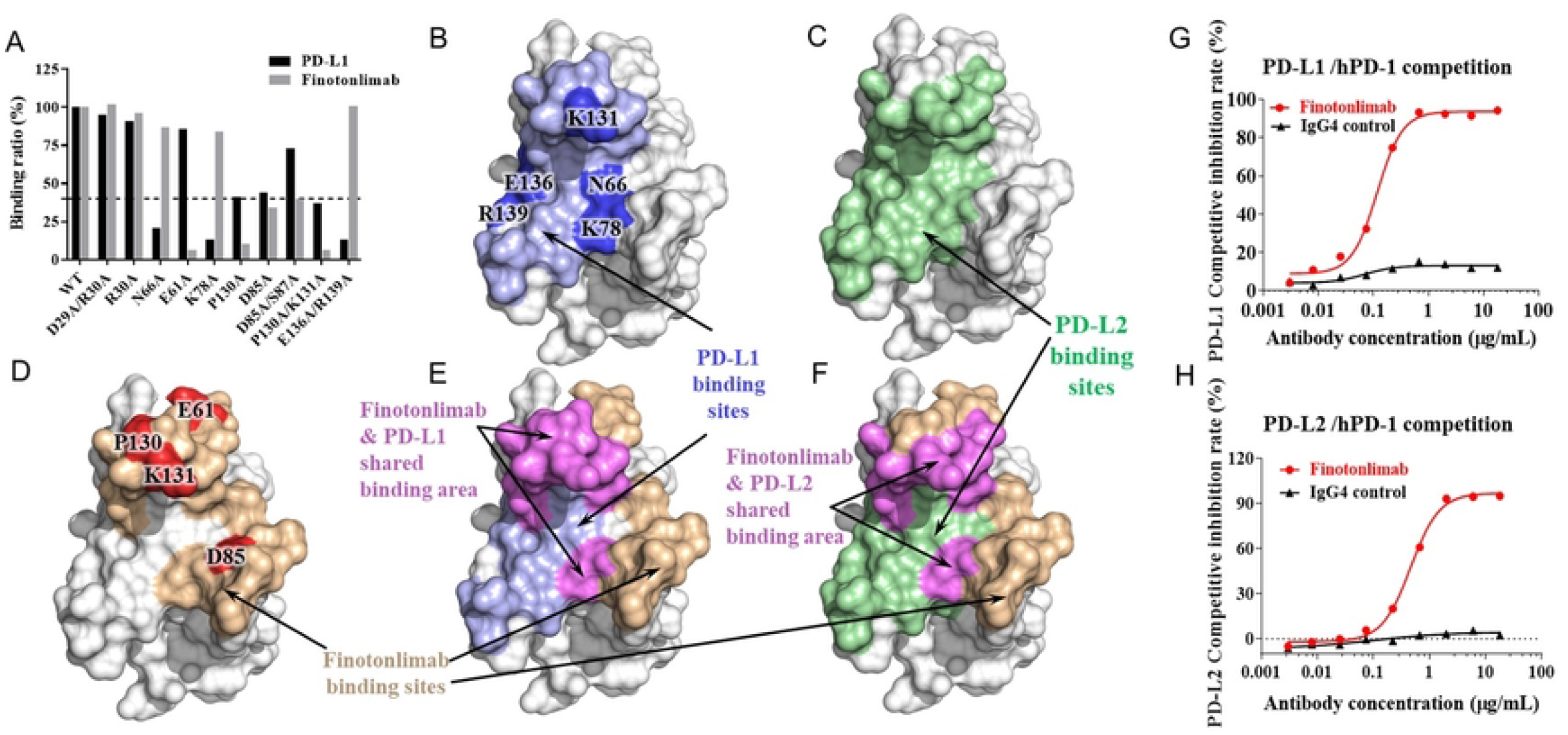
Finotonlimab blocks the binding of PD-1 ligands to PD-1and the epitope mapping of Finotonlimab. (A) The binding ratio of PD-L1/ Finotonlimab with PD-1 mutants (n=1). (B) PD-L1 binding sites in PD-1 are shown in light blue. The binding sites identified by mutation experiment are shown in blue. (C) PD-L2 binding sites in PD-1 are shown in green. (D) Epitopes of Finotonlimab identified by docking simulation and mutation experiment are shown in light red and red, respectively. (E) Binding sites of both PD-L1 and Finotonlimab in PD-1 are shown. The overlap of their binding sites is shown in magenta. (F) Binding sites of both PD-L2 and Finotonlimab in PD-1 are shown, the overlap of their binding sites is shown in magenta. Competitive inhibition rates of Finotonlimab to PD-L1/hPD-1 and PD-L2/hPD-1 are shown in (G) and (H), respectively.

Using those PD-1 mutants, the binding sites of Finotonlimab identified by the mutation experiment included E61, D85, and P130/K131 (**Figure 2D**). The complex structure of PD-1 and Finotonlimab was then constructed by docking the homology model of Finotonlimab with the PD-1 structure by using Discovery Studio, and taking the identified binding sites into consideration. The epitope of Finotonlimab in its complex structure with PD-1 largely overlaps with the binding sites of PD-L1, occupying about 44% of the PD-L1 binding area (**Figure 2E**), indicating that Finotonlimab effectively compete with PD-L1 for binding to PD-1. Similarly, Finotonlimab can also effectively competed with PD-L2 for binding to PD-1, occupying about 32% of the PD-L2 binding area (**Figure 2F**).

Compared with IgG4 isotype control, Finotonlimab efficiently blocked PD-1/PD-L1 and PD-1/PD-L2 binding, with calculated half-maximal inhibitory concentration (IC_50_) values of 0.116 and 0.462 μg/mL, respectively (**Figure 2G, 2H**). In a cell-based functional competition assay, Finotonlimab potently inhibited the binding of PD-L1 with PD-1 on Jurkat cells with an IC_50_ value of approximately 1.8 μg/mL compared with that of Nivolumab 2.5 μg/mL (**Figure S1**).

### *In vitro* Effects of Finotonlimab on the Function of T-cell Activation

The effects of Finotonlimab on T-cell function were assessed by T-cell reporter assay and mixed lymphocyte reaction (MLR). A PD-1/PD-L1 blocking bioassay system consisting of a Jurkat cell line stably transduced with human PD-1 and a nuclear factor of activated T-cell (NFAT)-driven luciferase reporter gene (Jurkat-PD-1-NFAT-Luc), and a CHO-K1 cell line expressing human PD-L1 and an artificial cell-surface TCR activator (CHOK1-PD-L1-TCRa) were utilized to demonstrate the T-cell activation effect. As shown in **Figure 3A**, the functional T-cell activation of Finotonlimab was determined by quantifying the luciferase signal level of reporter cell. The results suggested that Finotonlimab had a more potent concentration-dependent T-cell response activity as compared with Nivolumab.

**Figure 3.**
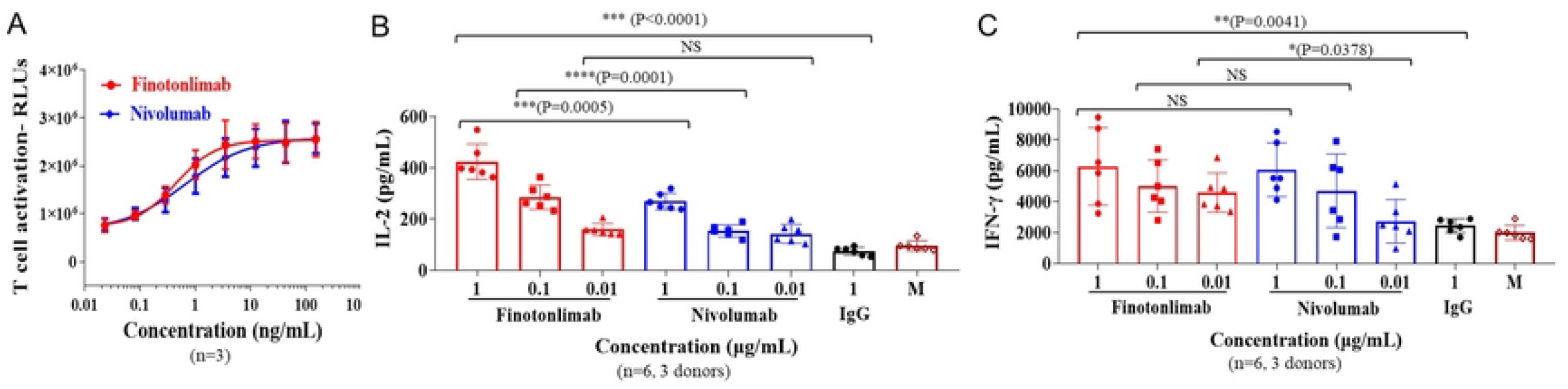
The functional activity of Finotonlimab in T-cell based assays. (A) T cell activation was determined by quantifying the luciferase signal level of reporter cell Jurkat-NFAT-Luc2p-PD-1 (n=3). The assay was evaluated by the interaction between effector cell Jurkat-NFAT-Luc2p-PD-1 and target cell CHO-K1-PD-L1-CD3E. In MLR assay, IL-2 (B) and IFN-γ (C) levels were determined using ELISA (CD4^+^ T cells and DC from the PBMCs of different donors were incubated in the presence of anti-PD-1 antibodies or isotype control at the indicated concentrations for 5 days, the PBMCs were isolated from 3 donors).

We then evaluated the functional antagonist activity of Finotonlimab in augmenting the response of primary human CD4^+^ T-cell in a T-cell MLR assay *in vitro*. Finotonlimab was shown to result in an increased T-cell activation, as measured by increased interleukin (IL)-2 and IFN-γ production in this assay. The enhanced activation of CD4^+^ T-cell induced by anti-PD-1 antibodies were concentration dependent (**Figure 3B, 3C**). It was showed that IL-2 level stimulated by Finotonlimab was significantly higher than that of Nivolumab at the same dose level. The levels of IL-2 stimulated by Finotonlimab and Nivolumab at the concentrations of 0.01, 0.1, 1 μg/mL were 159 ± 24, 286 ± 48, 424 ± 69 and 143 ± 37, 155 ± 24, 269 ± 32 pg/mL, respectively. The folds between the IL-2 level of Finotonlimab and Nivolumab at three concentrations were 1.11 (0.01 μg/mL), 1.84 (0.1 μg/mL) and 1.57 (1 μg/mL) (**Figure 3B**). The IFN-γ level stimulated by Finotonlimab was higher than that of Nivolumab. The IFN-γ levels stimulated by Finotonlimab and Nivolumab at the concentrations of 0.01, 0.1, 1 μg/mL were 4592 ± 1272, 5011 ± 1694, 6290 ± 2505 and 2742 ± 1402, 4707 ± 2375, 6070 ± 1728 pg/mL, respectively, while the differences were not significant at the concentration of 0.1 and 1 μg/Ml (**Figure 3C**).

### Finotonlimab Displayed Anti-tumoral Effects in Mouse Xenograft Models

The antitumor efficacy of Finotonlimab was investigated in a PD-1 humanized mouse model of MC38 tumors (**Figure 4A, 4B**). Tumor-bearing mice were treated with vehicle and Finotonlimab (2 and 8 mg/kg) every 3 days, successive treated for 6 times. Tumor volumes were measured twice a week until day 29 post first treatment. Relative tumor volume of Finotonlimab treated group compare to vehicle group indicate its tumor growth inhibition ability. The average tumor volume in vehicle group reached about 3405 ± 624 mm^3^ on the 28^th^ day, and the treatment with Finotonlimab at 2 and 8 mg/kg induced significant tumor growth suppression, the tumor volumes were reduced to 626 ± 150 and 484 ± 82 mm^3^, respectively. The tumor growth inhibitions (TGI%) were 85.3% and 89.7% at 2 mg/kg and 8 mg/kg, respectively, and 2 mg/kg group nearly reached optimal anti-tumor efficiency after infusion.

**Figure 4.**
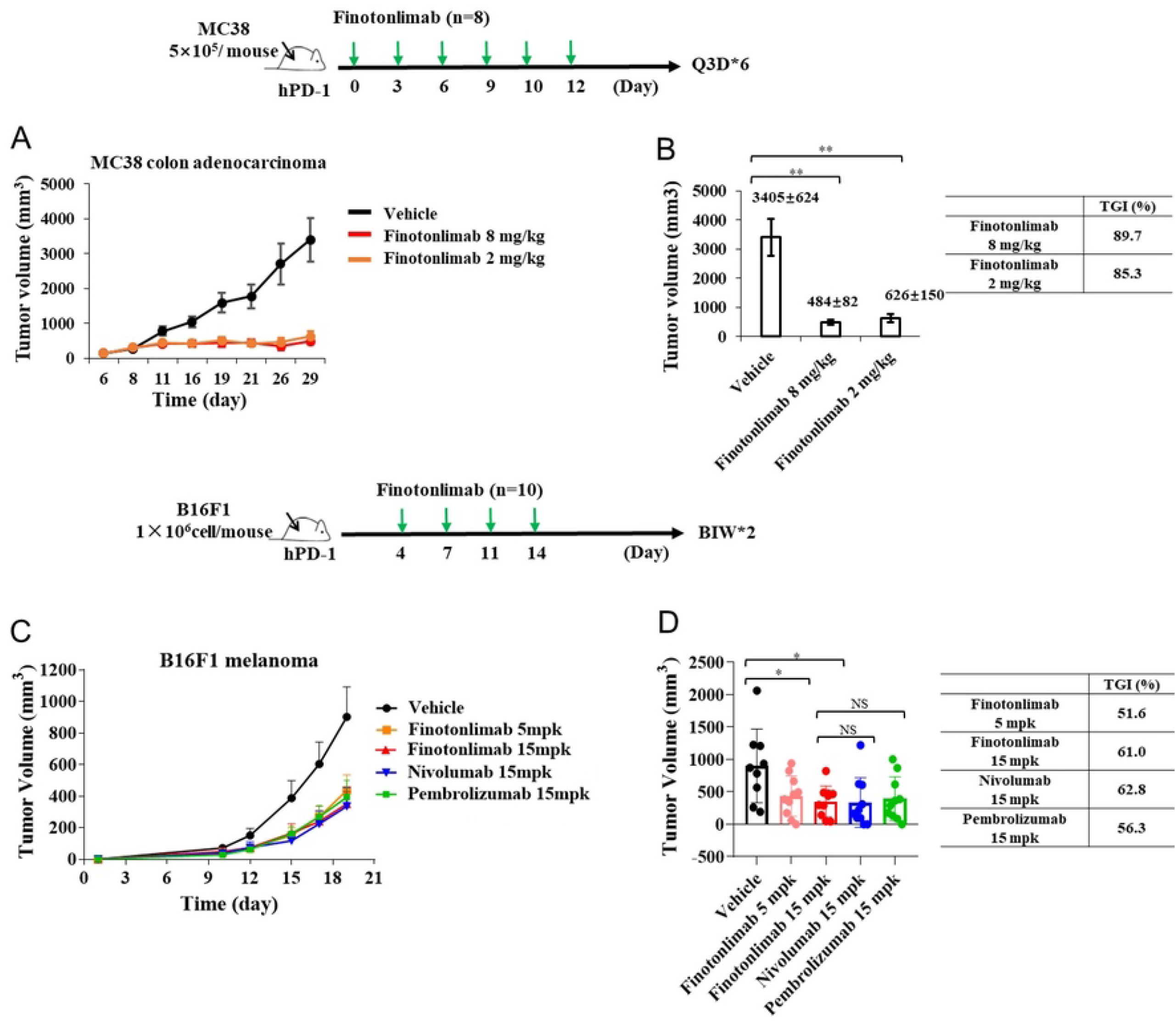
Antitumor activity of Finotonlimab monotherapy was evaluated in hPD1 humanized mouse models. The effects of Finotonlimab on mouse tumor volume (A) and percentage of tumor growth inhibition (TGI%) (B) in humanized mice with subcutaneous MC38 colon adenocarcinoma. The dosing schedule was every three days (Q3D) *6 (n=8 /group). The effects of Finotonlimab on mouse tumor volume (C) and TGI% (D) in humanized mouse model of B16F1 melanoma. The dosing schedule was twice a week (BIW) (n=10/group).

Next, we head-to-head evaluated the tumor control activity of Finotonlimab, Pembrolizumab, and Nivolumab using a PD-1 humanized B16F1 melanoma mouse model (**Figure 4C, 4D**). The results showed that the tumor growth was significantly inhibited by Finotonlimab, and the tumor volumes of Finotonlimab (15 mg/kg), Nivolumab (15 mg/kg), and Pembrolizumab (15 mg/kg) treated groups were comparable at day 19 (P>0.05). Moreover, the tumor growth inhibition (TGI%) of Finotonlimab treatment in the 15 mg/kg group (61.0%) was better compared to 5 mg/kg group (51.6%) (P>0.05). In addition to the hPD-1 humanized model, an hPBMC reconstructed human A431cutaneous squamous cell carcinoma tumor M-NSG mouse model was used to evaluate the anti-tumor efficacy of Finotonlimab. Mice with homogeneous tumor volume of A431 cells were treated with vehicle or Finotonlimab (10 mg/kg). Finotonlimab treated group showed slower tumor growth compared with vehicle group during the treatment, and at day 21, the tumor volumes of Finotonlimab treated and vehicle group were 680 ± 185 and 1362 ± 195 mm^3^, respectively, and TGI% was reached at 50% (**Figure 5A, 5B**).

**Figure 5.**
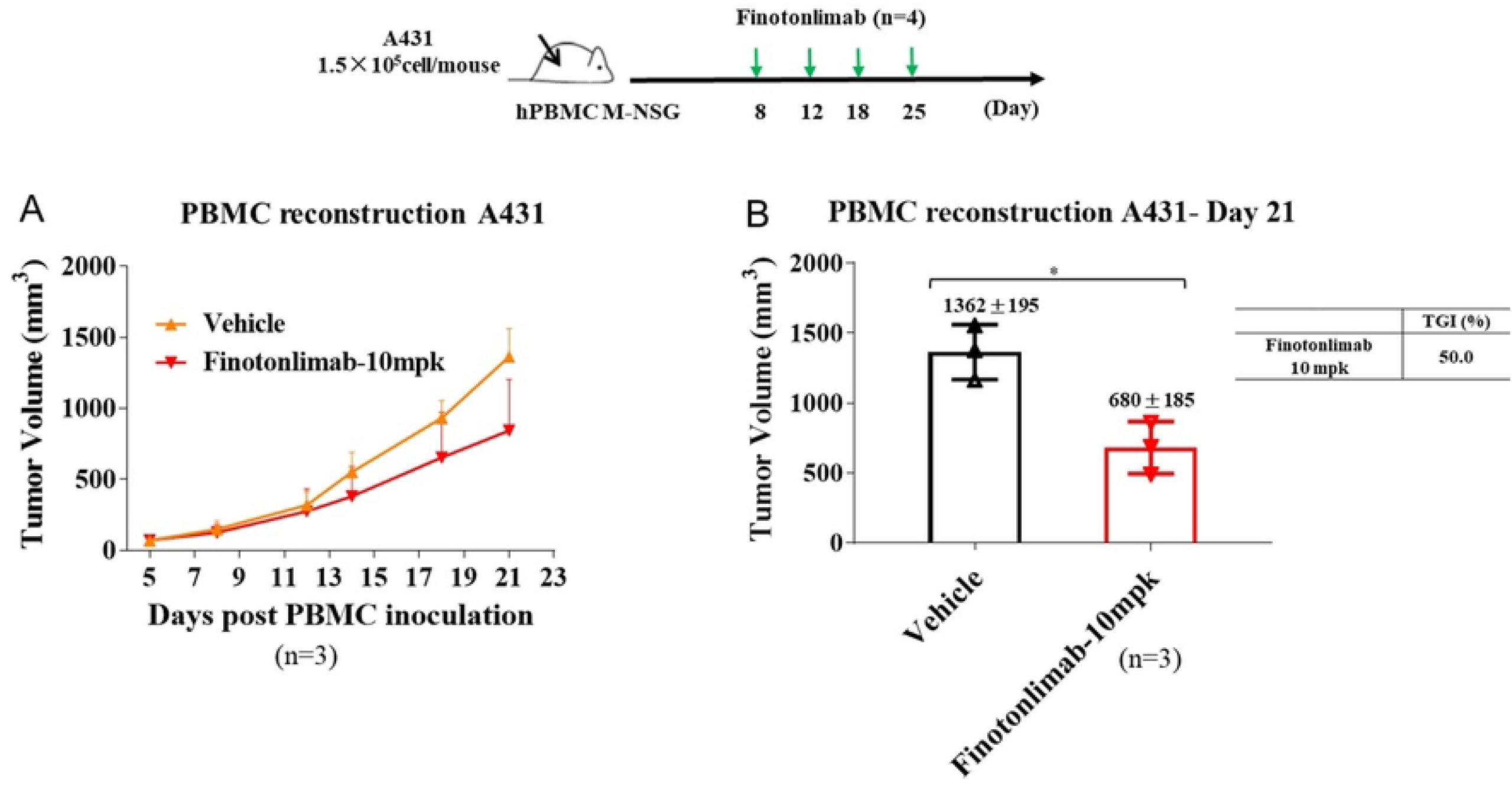
Antitumor activity of Finotonlimab monotherapy was evaluated in PBMC humanized tumor model. The effects of Finotonlimab on mouse tumor volume (A-B) in hPBMC humanized mice (A431 human epidermoid carcinoma).

### Fc Domain Mediated Functions of Finotonlimab

Immunoglobulins bound to cell surface receptors, such as PD-1, can attract natural killer (NK) cells, macrophages, and monocytes. Finotonlimab are of the human IgG4 subclass which has relatively low binding affinities to human FcγR, but retains binding to the human activating FcγRI receptor (CD64) [17]. The major function of FcγRI is to activate IgG-bound target cells via ADCP. FcγRIIIa (CD16a) is the primary receptor for NK- and macrophage-mediated ADCC. In this study, we detected the luciferase signals induced by the interactions between effector cells and target cells to evaluate the analog signals of Fc receptor related cytotoxic effect. The binding activity of anti-PD-1 antibodies with CD64, CD16a reconstructed cells and C1q protein was detected by FACS and ELISA, respectively. As expected, Finotonlimab induced positive but lower activation effects of the effector cell (Jurkat-NFAT-Luc2p-CD64), and the binding activity of Finotonlimab to FcγRI reconstructed cells was lower than those of Pembrolizumab (**Figure 6A, Figure S2A**). However, the effector cells (Jurkat-NFAT-Luc2p-CD16a) showed no luciferase signals on activated PD-1^+^ T cells in the presence of Finotonlimab **(Figure 6B)**, which is significantly lower than that of Pembrolizumab. Moreover, the lower binding activity of Finotonlimab with Jurkat-CD16A (F158) compared to that of Pembrolizumab was also observed (**Figure S2B**). Additionally, both results of complement dependent cytotoxic assay and normal human serum complements binding assay showed negative reaction response (**Figure 6C, Figure S2C**), suggesting that Finotonlimab had no evident CDC effect. These results indicated that Finotonlimab showed a similar Fc function of IgG4 isotype which minimize the negative influence on activated T cell and maintain excellent PD *in vivo*.

**Figure 6.**
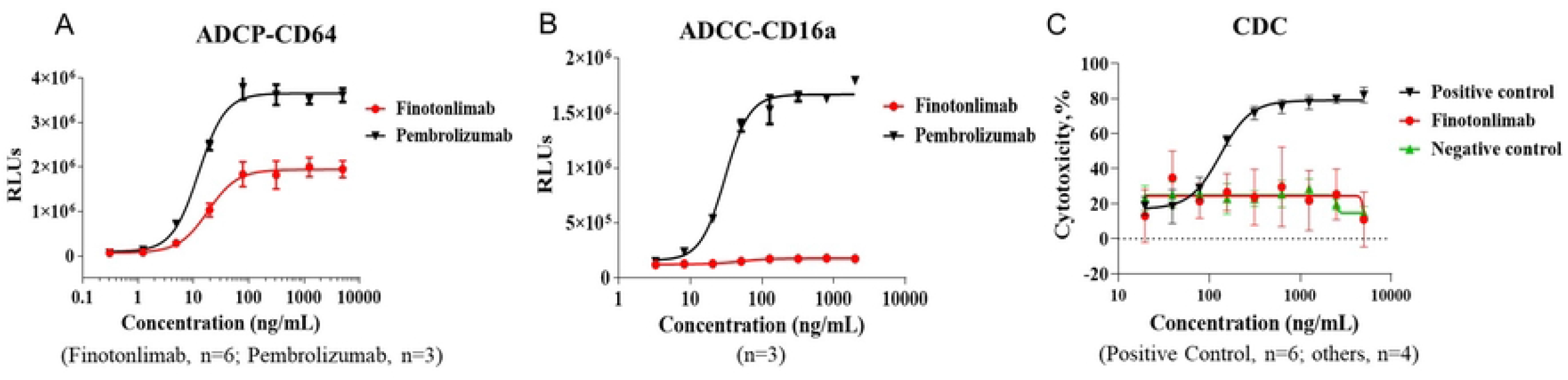
The analog signals of antibody-dependent cellular phagocytosis (ADCP), antibody-dependent cell-mediated cytotoxicity (ADCC), and complement-dependent cytotoxicity (CDC) of anti-PD-1 antibodies. (A) CD64 dependent ADCP of Finotonlimab and Pembrolizumab (n=6 and 3, respectively). (B) CD16a dependent ADCC of Finotonlimab and Pembrolizumab (n=3/group). (C) CDC of Finotonlimab and positive and negative control antibodies (n=4, 6, and 4, respectively).

### Pharmacokinetics Comparison of Finotonlimab and Anti-PD-1 Antibodies in Cynomolgus Macaques

It was suggested that pharmacokinetics correlated with pharmacology and efficacy [18, 19], herein the pharmacokinetics of Finotonlimab, Pembrolizumab, and Nivolumab were investigated in cynomolgus macaques. Following a single intravenous (i.v.) administration of anti-PD-1 mAbs at 5 mg/kg in cynomolgus macaques, standard pharmacokinetic (PK) measurements of Finotonlimab, Pembrolizumab and Nivolumab serum concentrations indicated a half-life (T_1/2_) of 206 h, 151 h and 198 h, respectively (**Figure 7A**). The C_max_ level of Finotonlimab was significantly higher than those of Pembrolizumab and Nivolumab.

**Figure 7.**
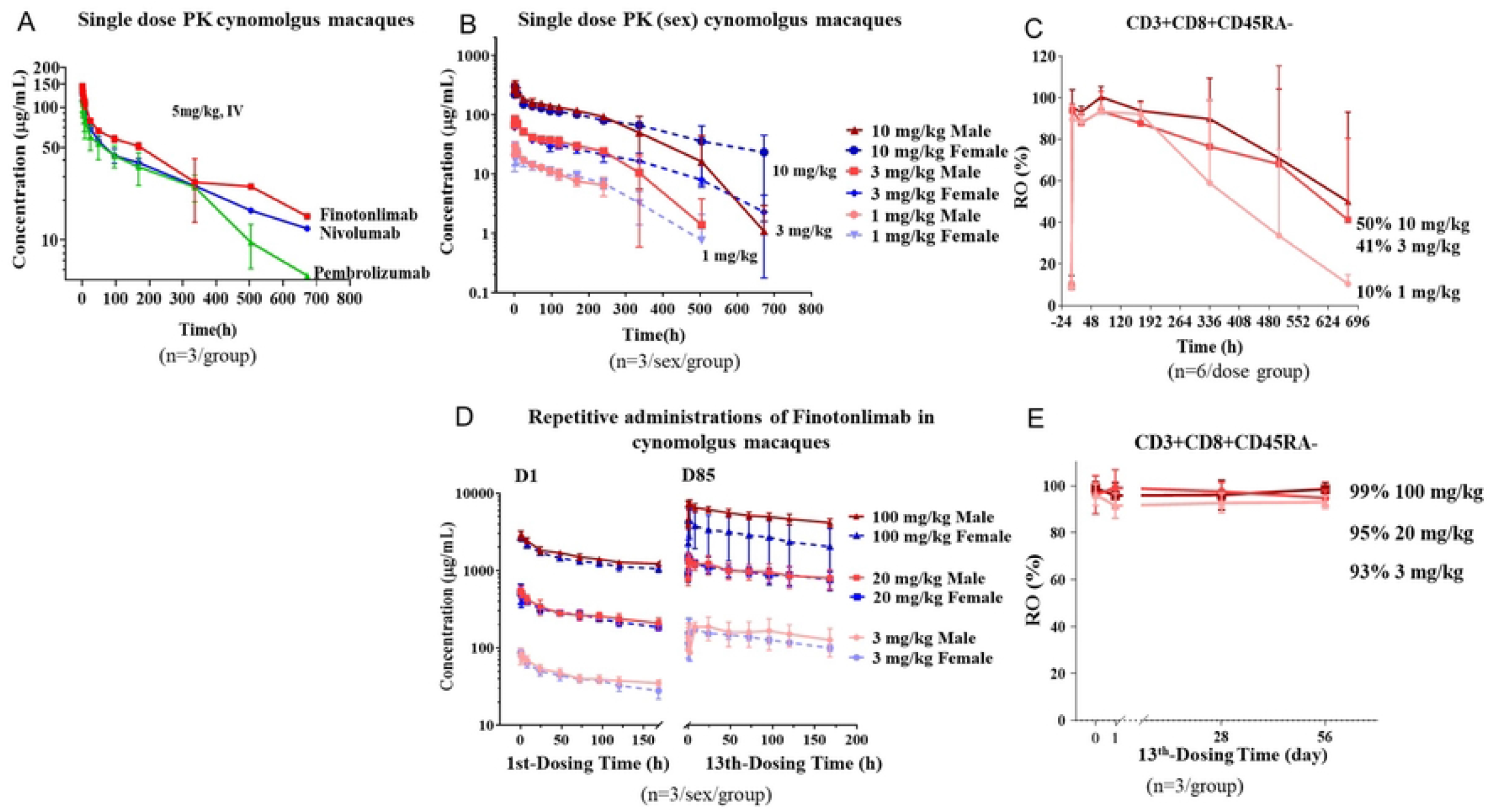
PD-1 receptor occupancy and Pharmacokinetic profiles. (A) Concentration-time profiles in serum after single intravenous administration of Finotonlimab, Nivolumab, or Pembrolizumab to cynomolgus macaques (n=3/group). (B) Concentration-time profiles in serum after single intravenous administration of different dose of Finotonlimab to cynomolgus macaques (n=3/sex/group). (C) PD-1 receptor occupancy profiles on CD3+CD8+CD45RA-cells after single administration of different doses of Finotonlimab. (D) Concentration-time curves in serum after repetitive administrations of Finotonlimab to cynomolgus macaques (QWx13). (E) PD-1 receptor occupancy profiles after the last administration (13^th^) of different doses of Finotonlimab.

### Pharmacokinetics and Pharmacodynamics of Finotonlimab in Cynomolgus Macaques by Single and Successive Administrations

Following a single intravenous (i.v.) administration of different doses of Finotonlimab (1, 3, 10 mg/kg) in cynomolgus macaques, the mean ± SD serum concentration-time curves of all dose treatments were shown according to different genders in **Figure 7B**. With the increase of dose from 1 to 10 mg/kg, the increased ratio of systemic exposure of Finotonlimab (AUC_inf_ and C_0_) was higher than the increased proportion of the corresponding dosages (**Table 2**). The concentration-time curves of Finotonlimab showed no gender differences in the first 120 hours at the same dose, while the differences were gradually obvious later which may be due to the production of anti-drug antibodies (ADA). In 3 and 10 mg/kg groups, all cynomolgus macaques had been determined ADA positive at days 29 post intravenous (i.v.) administration (**Table S1**). After a single intravenous injection of Finotonlimab, the PD-1 receptor occupancy (RO) rates of drug on non-naïve CD8^+^ T cells were saturated in all animals of all groups, and a plateau was also observed from 2 to 168 h (**Figure 7C, Table 3**). Then, a constantly slow decrease in RO% has been observed with dose-dependent relationship. At 672 h, the mean RO% in groups of 1, 3, and 10 mg/kg were 10.43% (range from 6.19% to 16.75%, n=6), 41.25% (range from 8.31% to 92.83%, n=6), and 50% (range from 8.3% to 95.38%, n=6), respectively.

**Table 2.** PK parameters of Finotonlimab after single intravenous administrations of 1, 3, 10 mg/kg to cynomolgus macaques (n=3/sex/group).

| Dose<br>(mg/kg) | Sex | / | T<sub>1/2</sub><br>h | C<sub>max</sub><br>µg/mL | AUC<sub>last</sub><br>h*mg/mL | AUC<sub>inf</sub><br>h*mg/mL | MRT<br>h | C<sub>max</sub><br>Ratio | AUC<br>Ratio |
| --- | --- | --- | --- | --- | --- | --- | --- | --- | --- |
| 1 | M | Mean | 178.34 | 30.13 | 2.71 | 4.46 | 92.09 | 1.00 | 1.00 |
|  |  | SD | 45.79 | 2.59 | 0.37 | 1.25 | 6.06 |  |  |
|  | F | Mean | 128.50 | 25.02 | 3.44 | 3.87 | 136.68 | 1.00 | 1.00 |
|  |  | SD | 25.43 | 9.95 | 0.85 | 0.91 | 25.12 |  |  |
| 3 | M | Mean | 169.15 | 84.58 | 10.89 | 14.40 | 132.52 | 2.81 | 4.02 |
|  |  | SD | 26.36 | 7.20 | 2.30 | 1.58 | 34.03 |  |  |
|  | F | Mean | 202.20 | 84.28 | 12.53 | 14.23 | 194.32 | 3.37 | 3.64 |
|  |  | SD | 41.68 | 5.71 | 1.23 | 2.47 | 6.66 |  |  |
| 10 | M | Mean | 191.71 | 339.39 | 44.22 | 61.49 | 151.24 | 11.26 | 16.33 |
|  |  | SD | 44.67 | 33.15 | 10.76 | 5.38 | 62.38 |  |  |
|  | F | Mean | 215.62 | 299.93 | 49.72 | 60.24 | 217.85 | 11.99 | 14.45 |
|  |  | SD | 154.11 | 27.57 | 10.02 | 20.85 | 48.02 |  |  |

**Table 3.** PD-1 receptor occupancy of different doses of Finotonlimab (n=6/group).

| Receptor occupation | 1 mg/kg |  | 3 mg/kg |  | 10 mg/kg |  |
| --- | --- | --- | --- | --- | --- | --- |
| Time for blood collection | Mean (%) | SD (%) | Mean (%) | SD (%) | Mean (%) | SD (%) |
| Before dose | 8.65 | 2.03 | 8.94 | 2.46 | 9.67 | 4.76 |
| 2h | 89.52 | 6.60 | 93.90 | 3.11 | 95.35 | 8.51 |
| 24h | 88.47 | 4.38 | 87.98 | 3.87 | 93.17 | 2.61 |
| 72h | 93.33 | 2.79 | 93.52 | 9.54 | 100.36 | 5.07 |
| 168h | 92.09 | 5.44 | 87.68 | 4.65 | 93.69 | 4.63 |
| 336h | 59.00 | 40.16 | 76.51 | 22.26 | 89.80 | 19.64 |
| 504h | 33.68 | 41.48 | 68.09 | 47.21 | 70.91 | 33.26 |
| 672h | 10.43 | 4.32 | 41.26 | 39.19 | 50.00 | 43.11 |

In the successive administration study, the serum drug concentrations of Finotonlimab for three doses (3, 20, and 100 mg/kg) were analyzed according to different gender. The maximum serum concentrations of Fintonlimab showed a dose dependent relationship (**Table S2, Figure 7D**). An obvious accumulation of Finotonlimab was observed in all groups. The AUC of final administration higher than that of first administration, with the accumulation factors (AI) (AUC_last_ (0-168 h) from day 85 and AUC_last_ (0-168 h) from day 1) of 1.45, 3.51 and 3.33 in males and 2.63, 3.63 and 2.03 in females, respectively (**Table S2**). The C_max_, AUC_last_, and AUC_inf_ in males were higher than those in female, the differences were consistent with the pharmacokinetics data in cynomolgus macaques (**Figure 7B**). After the 13^th^ administration, the Finotonlimab RO performance saturated in cynomolgus macaques in all dose groups (**Figure 7E**), and a sustained PD-1 receptor occupancy of more than 93% lasted up to 8 weeks. The high receptor occupancy rate of Finotonlimab in successive administrations was consistent with that in a single administration, suggesting that even a low dose of Finotonlimab treatment could sustained the receptor occupancy to a saturated level.

### Toxicology Evaluation by Successive Administrations of Finotonlimab in Cynomolgus Macaques

The toxicology of Finotonlimab was further evaluated in cynomolgus macaques. After successive intravenous dosing once every week for 13 weeks, Finotonlimab treatment was well tolerated in cynomolgus monkeys with no mortality or morbidity noted in any groups at doses of 3, 20, or 100 mg/kg. There also no obvious treatment related abnormal changes in clinical observations, body weight, body temperature, ECG (Electrocardiograph), blood pressure, ophthalmoscopic examinations, coagulation function, blood biochemistry, urinalysis analysis and organ weights in any groups. Inflammatory cell infiltration noted in multiple organs was related to pharmacological function and reversed after a recovery period up to 8 weeks. Therefore, it is considered that the NOAEL (no observed adverse effect level) of Finotonlimab is more than or equal to 100 mg/kg.

## Discussion

This study introduced functional characterization of Finotonlimab, a humanized anti-PD-1 monoclonal IgG4, including the PD-1 affinity, ligands blocking, *in vitro* T cell activation effect and *in vivo* anti-tumor activity in xenograft naïve mice or hPBMC reconstructed mice, Fc-mediated effector functions, RO, PK and safety characters of Finotonlimab after single dose or successive doses in cynomolgus macaques. Those data indicated that Finotonlimab exhibits an effective and safe anti-PD-1 profile, and its anti-tumor efficacy will be evaluated and verified in several solid tumor types in Phase III trials.

Finotonlimab binds specifically to human PD-1 receptor with a high binding affinity (K_D_) of 64.8 pM, and antagonizes the interaction of PD-1 with its ligands PD-L1 and PD-L2, which resulted in the potent activation of T cell activity, represented by significantly enhanced IL-2 and IFN-γ production with a stimulatory effect on CD4^+^ T cells in MLR assay. Finotonlimab exhibited potent anti-tumor efficacy in A431 xenograft hPBMC reconstructed mice, which furthest mimic the distinct features of the corresponding human immune system and tumor microenvironment. In another immune competent mouse model of B16F1 melanoma, 15 mg/kg Finotonlimab treatment lead to a strong anti-tumor effect against B16F1 tumors which is comparable with Nivolumab and Pembrolizumab. The findings of these studies indicated a promising preclinical evaluation of Finotonlimab as a checkpoint inhibitor and provided a basis for the potent anti-tumor efficacy in the ongoing clinical trials [20].

Another critical aspect in predicting the effector T cell functions and safety profile of a new anti-PD-1 therapy is cytotoxicity induced by the binding to complement of Fcγ receptors [21, 22]. Evidences have demonstrated that anti-PD-1 antibodies are needed to lower Fc-mediated effector functions (ADCC, ADCP, and CDC) to avoid the killing of PD-1^+^ T cells by FcγR^+^ effector cells [17, 23]. In our studies, the Fc-mediated effector functions were demonstrated by analog signals of ADCC and ADCP using the recombinant reporter cells. Consistent with its IgG4 framework, Finotonlimab showed weak FcγRIII (CD16) induced ADCC or CDC associated cytotoxicity [17, 21, 24]. In addition, comparative characterization data demonstrated that Finotonlimab induced lower activated FcγRI (CD64) mediated ADCP activity and FcγRIII (CD16) induced ADCC activity than Pembrolizumab, and hence Finotonlimab is unlikely to result in the high depletion capacity of antitumor effector T cell [23, 25-28].

As mentioned before, anti-PD-1 mAbs bind to PD-1 receptors, blocking immune-suppressing ligands from interacting with PD-1, thus restoring T-cell and immune responses [29-31]. Therefore, high percentage and sustained PD-1 receptor occupancy are likely to result in a better anti-tumor effect and a stronger immune response [30]. After single intravenous administration of Finotonlimab to cynomolgus macaques, the results showed that the drug occupied hPD-1 on T-cell to a saturated state and sustained a slow off-rate for a long time along with decreasing serum drug concentration, potentially giving a longer duration of drug effect. After successive administrations of different doses of Finotonlimab, high level of PD-1 occupancy on circulating T cells by drug was observed consistently for up to 8 weeks post the last administration in each group. Of note, even 1 mg/kg of Finotonlimab injected to cynomolgus monkeys achieved more than 93% of receptor occupancy.

Furthermore, the possible PK advantage of anti-PD-1 antibodies could indicate better pharmacodynamics and drug efficacy [19, 30, 32]. Single dose PK study confirmed a better PK profile of Finotonlimab than that of Pembrolizumab and Nivolumab at the same dose level in cynomolgus macaques. After successive doses of Finotonlimab in cynomolgus macaques, the increasing ratio of drug exposure (AUC_0-last_ and C_0_) in all groups was higher than that of dose, as shown in **Table S2**. The same trend was observed in single dose PK study of Finotonlimab in **Table 2**. AUC _(0-168h)_ of last administration was higher than that of the first administration with obvious drug accumulation (**Table S2**). Sustainable high level of PK-RO profile of Finotonlimab in animals provided the basis for the lasting pharmacodynamics and stronger immune response in humans.

Immunogenicity evaluation is a critical detection index during clinical development of antibody-based drugs. However, in single dose PK study, the clearances of 3 and 10 mg/kg groups in the later releasing period were more rapid than that of 1 mg/kg group, which presented non-linear profile (**Figure 7B**). It might be due to the production of ADA in middle and high dose groups. Indeed, ADA were detected in some animals in all groups (**Table S1, Table S3**). The incidence of ADAs for anti PD-1 antibodies have been reported frequently in several Food and Drug Administration approved products, while Finotonlimab showed similar characteristics as other anti-PD-1 drugs as well [33, 34]. However, the anti-tumor efficacy and safety of Finotonlimab needs to be further confirmed in clinical trials.

In conclusion, as a new humanized anti-PD-1 monoclonal antibody, Finotonlimab has a high affinity to PD-1 with lower Fc mediated function, and shows lasting PK/PD profiles and potent *in vivo* anti-tumor effects. The non-clinical data are encouraging and could provide a basis for its efficacy and safety evaluation in clinical trials.

## Materials and Methods

### Reagents

Finotonlimab was manufactured by SinoCellTech (Beijing, China). Nivolumab was purchased from Bristol-Myers Squibb (USA). Keytruda-biosimilar (anti-PD1(MK)-IgG4) was purchased from Sino Biological (Beijing, China). B7-H1-Fc (PD-L1) recombinant protein, PD-1 wt/mutant proteins, the secondary antibody of goat anti-Human IgG Fc/HRP, the negative control (H7N9-R1-IgG4) and the positive control (PD1-H944-1-IgG1(o), 20171009) were purchased from Sino Biological (Beijing, China). Goat anti-Human IgG F(ab)2 /HRP was purchased from JACKSON (USA). The recombinant proteins of PD-1-his, CD28-Fc, CTLA4-His, BTLA-Fc, PIGF-Fc and Kappa-R002/HRP were purchased from Sino Biological (Beijing, China). B7-H1-His-biotin, PD-L2-His-biotin, and PD-1-Fc were purchased from Sino Biological (Beijing, China). Streptavidin/HRP was purchased from ZSGB-BIO (China). Streptavidin Alexa Fluor® 488 Conjugate was purchased from Life Technologies. The secondary antibody of PE Mouse Anti-Human CD279 was purchased from BD Biosciences. rhGM-CSF (215-GM-010) was purchased from R&D. rhIL-4 (GMP-11846-HNAE) was purchased from Sino Biological. Human IFN-γ ELISA Set (555142) and Human IL-2 ELISA Set (555190) were purchased from BD OptEIA™. CD4 microbeads (130-045-101), anti-FITC microbeads (130-148-701) and LS Columns (130-042-401) were purchased from Miltenyi Biotec. FITC Mouse Anti-Human CD4 (555346) was purchased from BD Biosciences. Ficoll-PaqueTM PLUS (17144002) was purchased from GE. Luciferase Assay System (E1501) and Passive Lysis 5 × Buffer (E1941) were purchased from Promega. C1q recombinant protein (A400) was purchased from Quidel and its secondary anti C1q/HRP (ab46191) from Abcam.

### Animals

Human PD-1 knock-in C57BL/6 mice (B-hPD-1/C57BL/6) were purchased from Biocytogen Pharmaceuticals (Beijing) Co., Ltd. All animal experiments were performed in accordance with regulations for care and use of laboratory animals at Biocytogen Pharmaceuticals, and were approved by Institutional Animal Care and Use Committee. All mice were kept in specific pathogen-free conditions.

### Affinity measurements by Octet

Recombinant human PD-1 protein was biotinylated and loaded using SA sensor (Pall Corporation). Six concentrations of Finotonlimab and Nivolumab were added for real-time association and dissociation analysis using Octet system. Data Analysis Octet was used for data processing.

### Epitope mapping of PD-L1/PD-L2 binding

96-well plates were coated (100 μL/well) with PD-1 wt/mutant proteins diluted to 1 μg/mL at 4°C overnight. After washing, the plates were blocked for over 1 h at room temperature. Then PD-L1-Fc/ PD-L2-Fc was diluted to 10 μg/mL and added 100 μL/well for 2h. After washing out the free antibodies, 250 ng/ml IgG Fc/HRP was added for 70 μL/well, then incubated for 1 h at room temperature and washed the wells. Finally, the assay was conducting chromogenic reaction followed by absorbance measurement (450 nm) in an ELISA reader.

### Epitope mapping of Finotonlimab binding

96-well plates were coated (100 μL/well) with PD-1 wt/mutant proteins diluted to 10 ng/ml at 4°C overnight. After washing, the plates were blocked for over 1 h at room temperature. Then Finotonlimab or negative antibodies were diluted to 2 μg/mL, added 100 μL/well and incubated for 2 h at room temperature. After washing out the free antibodies, 250 ng/mL second antibody of IgG Fc/HRP was added for 70 μL/well, then incubated for 1 h at room temperature and washed the wells. Finally, the assay was conducting chromogenic reaction followed by absorbance measurement (450 nm) in an ELISA reader.

### Structural modeling of Finotonlimab/PD-1 complex and epitope analysis

The structure of Finotonlimab was firstly homology modeled by Discovery Studio 4.0 (Accelrys Software). The structure of PD-1 was extracted from the crystal structure (PDB ID 5GGS). Then the complex structure of PD-1 and Finotonlimab was constructed by docking of the structures of Finotonlimab and PD-1 in Discovery Studio 4.0 according to the binding sites that identified by the mutation experiment. The docked complex structures were optimized by energy minimization and the complex structure with the lowest free energy was selected. For the binding interface analysis, the residues of PD-1within 5.0 Å of Finotonlimab (modeled complex structure), PD-L1 (PDB ID 4ZQK), or PD-L2 (PDB ID 6UMT) were recognized as the binding area of Finotonlimab, PD-L1, or PD-L2, respectively, and displayed by PyMOL Molecular Graphics System, Version 2.4 (Schrödinger, LLC).

### Binding affinity of Finotonlimab to human PD-1 by ELISA

The affinity of the antibody to human PD-1 protein was assessed by ELISA. 96-well plates were pre-coated (100 μL/well) with different concentration of human PD-1 protein (0.16, 0.49, 1.48, 4.44, 13.33, 40, 120, 360, 1080, 3240 and 9720 ng/mL) at 4°C overnight. After washing, the plates were blocked for over 1 h at room temperature, then Finotonlimab, Nivolumab and H7N9-R1-IgG4 negative control were added for 2 μg/ml (100 μL/well) and were incubated for 2 h at room temperature. After incubation and rinsing, HRP-labeled goat anti-human IgG antibody was added, incubated at room temperature for 1 h, rinsed, and developed with TMB substrate for 10 min. The reaction was stopped with 2 M HCl, and the absorbance was read at 450 nm using a microplate spectrophotometer within 15 min. Origin 7 software was used to calculate EC_50_ and draw the profiles.

### Binding of Finotonlimab to human PD-1 family proteins by ELISA

96-well Plates were pre-coated with different concentrations (1.25, 5, 20, 80, 320, 2560 and 10240 ng/mL) of human PD-1-His, CD28-Fc, CTLA4-His, BTLA-Fc or PIGF-Fc at 4°C overnight. After washing, the plates were blocked for over 1h at room temperature, then Finotonlimab, Nivolumab and H7N9-R1-IgG4 negative control were added for 2 μg/mL (100 μL/well) and were incubated for 2 h. After incubation and rinsing, HRP-labeled goat anti-human IgG antibody was added, incubated at room temperature for 1 h, rinsed, and developed with TMB substrate for 10 min. The reaction was stopped with 2 M HCl, and the absorbance was read at 450 nm using a microplate spectrophotometer within 15 min. Origin 7 software was used to calculate EC_50_ and draw the profiles.

### Binding of Finotonlimab to C1q by ELISA

96-well Plates were pre-coated with different concentrations of anti-PD-1 antibodies at 4°C overnight. After washes, the plates were blocked for over 1 h at room temperature, then C1q recombinant protein was added (5 μg/mL, 100 μL/well) and incubated at room temperature for 2 h. After removing the unbound C1q recombinant protein, secondary anti-C1q/HRP antibody was added (5 μg/mL, 100 μL/well), incubated at room temperature for 1 h, rinsed, and developed with TMB substrate for 10 min. The reaction was stopped with 2 M HCl, and the absorbance was read at 450 nm using a microplate spectrophotometer within 15 min. Origin 7 software was used to calculate EC_50_ and draw the profiles.

### Evaluating human PD-1/PD-L1 and PD-1/PD-L2 blocking capability of Finotonlimab by ELISA

ELISA was used to evaluate the competitive blocking capability of the testing antibody to PD-L1/PD-L2 against PD-1.

96-well Plates were pre-coated with human 0.4 μg/mL PD-1-mFc (100 μL/well), then incubated at 4°C overnight. After 1 h of blocking, 100 μL 1 μg/mL B7-H1-His and different concentration (0.003, 0.008, 0.025, 0.074, 0.222, 0.667, 2, 6, 18 μg/mL) of Finotonlimab or Nivolumab or H7N9-R1-IgG4 negative control was added, and were incubated at room temperature for 2 h, respectively. After incubation and rinsing, streptavidin/HRP was added, incubated at room temperature for 1 h, rinsed, and developed with TMB substrate for 10 min. The reaction was stopped using 2 M HCl, and the absorbance was read at 450 nm using a microplate spectrophotometer within 15 min.

For the assay of evaluating human PD-1/PD-L2 blocking capacity of Finotonlimab, the concentration of PD-1-mFc to pre-coat wells was 2 μg/mL. After 1 h of blocking, 100 μL 1 μg/mL PD-L2-His-biotin and different concentration (0.003, 0.008, 0.025, 0.074, 0.222, 0.667, 2, 6, 18 μg/mL) of Finotonlimab or Nivolumab or H7N9-R1-IgG4 negative control was added, respectively, and were incubated for 2 h. Other steps were same as above. The percentage of inhibition rate was calculated as PI% = (OD_blank_-OD_sample_)/OD_blank_×100%. OD_blank_ refers to negative control group, OD_sample_ refers to antibody group.

### Binding of the Finotonlimab to cell surface PD-1 by FACS

FACS was used to evaluate the competitive blocking capability of the testing antibody to PD-L1 against PD-1 on the cell surface. Jurkat/ PD-1 was incubated with anti-PD-1 antibodies or negative control (2-fold serially diluted from 200 μg/mL to 0.195 μg/mL) at 4°C for 20 min. After rinsing with PBS and centrifuged, a constant concentration of FITC labeled Goat anti-human IgG Fc was added and incubated at 4°C for 20 min. After washing out the unlabeled secondary antibody, 200 μL PBS was added to re-suspend cells and the resulted cell suspension was filtered with cell strainer. The MFI of the cells was measured by flow cytometry.

The methods of Finotonlimab cell binding assay to the surface of CD16a (Jurkat-NFAT-Luc2p-CD16A) and CD64 (Jurkat-NFAT-Luc2p-CD64) reconstructed cells were the same as above.

### Evaluating human PD-1/PD-L1 blocking capability of Finotonlimab by FACS

FACS was used to measure the affinity of the anti-PD-1 antibody to the cellular surface PD-1 protein. The CHO cells (an engineered cell line expressing the human PD-1 protein) were incubated with PD-1 antibodies or negative control (2-fold serially diluted from 10 μg/mL to 0.26 μg/mL) at 4°C for 20 min. After the cells were washed, 10 μL of B7H1-Fc-biotin (0.4 μg/mL) was added and washed to remove antibodies. Then the secondary antibody of streptavidin-Alexa Fluor® 488 conjugate was added and incubated at 4°C for 20 min. After rinsing and leaving out the free secondary antibodies, 200 μL PBS was added to re-suspend the cell. Finally, filtered the cell suspension with cell strainer and measured the MFI by flow cytometry. The percentage of inhibition rate was calculated as PI% = (MFI_blank_-MFI_Sample_)/MFI_blank_×100%. OD_blank_ refers to negative control group, OD_sample_ refers to antibody group.

### T cell activation associated luciferase reporter assays

The stimulation effect of anti–PD-1 treatment on Jurkat T cells was evaluated using Jurkat PD-1/PD-L1 reporter system, the details were shown as previously described [35]. Target cell CHO-K1-PD-L1-CD3E was seeded with 2 × 10^4^ cells/well in the 10% FBS DMEM culture medium overnight. After removing the supernatant, different concentrations of PD-1 antibodies were added. Then effector cell of Jurkat-NFAT-Luc2p-PD-1 with the concentration of 7.5×10^4^ cells/well was dispensed into CHO-K1 cells and the plates were incubated at 37°C and 5% CO_2_ for 6 h. After incubation, Passive Lysis 5 × Buffer with 20 μL/well was added and mixed well. Then 20 μL supernatant was transferred to a clean 96 well plate and the absorbance was read using a microplate spectrophotometer within 15 min.

### *In vitro* mixed lymphocyte reaction

CD4+ T cells were isolated from the PBMCs of a healthy human donor and dendritic cells were derived from a different donor, via isolation of monocytes. Following 3 days of culturing with rhGM-CSF (20 ng/mL) and rhIL-4 (160 ng/mL) in 10% FBS contained 1640 culture medium at 37°C, 5% CO2, half of the culture medium need to be changed every time. After 6 days, the dendritic cell suspensions were collected. Dendritic cells (1×10^4^ cells/well) and allogeneic CD4^+^ T cells (1×10^5^ cells/well) were incubated in the presence of various concentrations (0.01, 0.1, 1 μg/mL) of anti-PD-1 antibodies or isotype control antibody at 37°C, 5% CO2 for 5 days, and T-cell activation was quantified by the level of IL-2 and IFN-γ secreted in culture supernatants as determined by ELISA.

### ADCC/ADCP-activation assays

As previous studies described, we confirmed the FcγR functions by generating the reporter system [53]. For the ADCC (CD16a) or ADCP (CD64) assay, the target cells CHO-PD-1 were seeded on 96-well plates and were cultured in 10% FBS contained DMEM medium at 37°C, 5% CO2 overnight. After removing the supernatant, 0.5 g/L PF68 contained RPMI 1640 (Phenol Red-free) culture medium was used to wash the cells for 2 times. Then different concentrations of antibodies (40 μL/well) and 1×10^5^ cells/well effector cells Jurkat-NFAT-Luc2p-CD16a or Jurkat-NFAT-Luc2p-CD64 were added, cultured at 37°C, 5% CO2 for 4 h. After incubating, passive lysis 5 × Buffer with 20 μL/well was added and mixed well. Then 20 μL supernatant was transferred to a clean 96 well plate and the absorbance was read using a microplate spectrophotometer within 15 min.

### CDC cytotoxicity assay

96 well plates were seeded with CHO-PD-1 for 5×10^4^ cells/well in 0.1% BSA contained 1640 culture medium (Phenol Red-free). Then different concentrations of antibodies (50 μL/well) combined with the complement (50 μL/well) at the dilution rate of 1:4 were added to the wells. After incubation at 37°C for 3 h, 15 μL of WST-8 substrate solution was added and the absorbance was read using a microplate spectrophotometer within 15 min.

### Efficacy study in MC38 tumor mouse models

The mouse colon carcinoma MC38 cells were subcutaneously inoculated to the right flank of the hPD-1 knock-in mice (5 × 10^5^ cells/mouse). When the tumor size reached about 150 mm^3^, the mice were randomly divided into three groups: Vehicle group, 2 mg/kg Finotonlimab group, 8 mg/kg Finotonlimab group, with eight mice in each group. After treatment, the efficacy assessment was performed by evaluating the tumor volume and tumor growth inhibitory rate (TGI%).

### Efficacy study in B16F1 tumor mouse models

The mouse melanoma cells B16F1 cells were subcutaneously inoculated to the right back of the hPD-1 knock-in mice (1 × 10^6^ cells/mouse). When the tumor size reached about 150 mm^3^, the mice were randomly divided into five groups: Vehicle group, 5 mg/kg Finotonlimab group, 15 mg/kg Finotonlimab group, 15 mg/kg Pembrolizumab group and 15 mg/kg Nivolumab group with ten mice in each group. After treatment, the efficacy assessment was performed by evaluating the tumor volume and tumor growth inhibitory rate (TGI%).

### Efficacy study in hPBMC reconstituted A431 mouse models

A431 cell line was incubated RPMI 1640 medium supplemented with 10% fetal bovine serum. The cell suspension was harvested and adjusted to 1.5×10^6^ cell/mL and 25% matrigel was added. After anesthesia and shaving, each M-NSG mouse was implanted subcutaneously in the right back with 100 μL A431 (1.5 × 10^5^ cell/mouse). The next day, hPBMC was recovered with RPMI 1640 medium supplemented with 10% fetal bovine serum and transferred into the T75 cell culture flask. After rinsing with PBS and centrifuging at 200 × g for 15 min to collect cells, the cell suspension was adjusted to 5×10^7^ cells/ml in ice-cold PBS. The obtained hPBMC cell suspension was injected i.v. via tail vein (100 μL, 5 × 10^6^ cells).

### Pharmacokinetics and toxicity assessment of Finotonlimab in cynomolgus monkeys

In a single-dose pharmacokinetic study, cynomolgus monkeys received Finotonlimab at 1, 3, 10 mg/kg (3/sex/group) by i.v. administrations. Nine male and nine female cynomolgus monkeys were selected and randomly divided into the three dose groups. The blood samples for PK assessment were collected before (0) dosing and at 0.042, 1, 2, 8, 24, 48, 72, 96, 120, 168, 240, 336, 504 and 672 h after dosing, respectively.

In a 13-week toxicology study, cynomolgus monkeys received 13 successive weekly i.v. administrations of 3, 20 and 100 mg/kg of Finotonlimab, administered at a constant volume of 0.4 ml/kg, followed by an 8-week recovery period. After the last administration, three monkeys/sex of each group were euthanized and the remaining animals were euthanized at the end of recovery period.

### Receptor occupancy assay

The whole blood sample was collected at 2, 24, 72, 168, 336, 504 and 672 h post the first administration in accompany with PK assessment. The receptor occupancy assay was measured by FACS.

Lysing erythrocytes. 700 μL whole blood sample was collected and was incubated with 7 ml lysing buffer for 15 min at room temperature. Then erythrocytes were washed by PBS, centrifuged at 300 × g at 25°C for 5 min, and re-suspended in 300 μl of PBS. And the cell suspension was divided into four tubes (A, B, C, and D).

Staining and incubation. Anti-PD-1 antibodies or H7N9 antibody were added into the tubes of C and D and the mixture was vortexed, incubated at 4°C for 20 min. Then 3 mL of PBS was added into A, B, C, and D and the cell suspension was centrifuged at 1400 × rpm for 5 min and the supernatant was removed. 3 ml of PBS buffer was added to wash the cells. The cells were centrifuged at 1400 × rpm for 5 min with the supernatant removed. Human IgG antibody was added into the tubes of B, C, and D. The tubes were vortexed and incubated at 4°C for 15 min. All tubes were washed for 2 times with 3 mL of washing buffer and centrifuged at 1400 × rpm for 5 min with the supernatant removed. The antibodies of CD3, CD4, CD8 and CD45RA were then added with the mixture vortexed and incubated at 4°C for 20 min. Then the cells were washed for 2 times with 3 mL of washing buffer and centrifuged at 1400 × rpm for 5 min. After filtered the cell suspension with cell strainer, the receptor occupancy of all tubes was measured by FACS assay.

## Author Contributions

LZX and CYS designed the studies, CYS, XNY and JL collected the data and contributed to manuscript preparation. XZ, JLJ, RW and CLY performed the animal experiments. EHG and MJD performed the antibody experiments. JM, YQD, LLS and SL performed the in vitro experiment.

## Acknowledgments

Non.

## References

1. Lee JJ, Chu E. Recent Advances in the Clinical Development of Immune Checkpoint Blockade Therapy for Mismatch Repair Proficient (pMMR)/non-MSI-H Metastatic Colorectal Cancer. Clinical Colorectal Cancer. 2018:S1533002818302767-.

2. Centanni M, Moes D, Trocóniz I, Ciccolini J, Van Hasselt JGC. Clinical Pharmacokinetics and Pharmacodynamics of Immune Checkpoint Inhibitors. Clinical Pharmacokinetics. 2019.

3. Ishida, Agata, Shibahara, Honjo. Induced expression of PD-1, a novel member of the immunoglobulin gene superfamily, upon programmed cell death. The EMBO journal. 1992.

4. Yao S, Chen L. PD-1 as an immune modulatory receptor. Cancer J. 2014;20(4):262–4. doi: 10.1097/PPO.0000000000000060. PubMed PMID: 25098286; PubMed Central PMCID: PMCPMC4455017.

5. Nishimura H, Nose M, Hiai H, Minato N, ∣∣ TH. Development of Lupus-like Autoimmune Diseases by Disruption of the PD-1 Gene Encoding an ITIM Motif-Carrying Immunoreceptor. Immunity. 1999;11(2):141–51.

6. Yasutoshi A, Akemi K, Hiroyuki N, Yasumasa I, Takeshi T, Hideo Y, et al. Expression of the PD-1 antigen on the surface of stimulated mouse T and B lymphocytes. International Immunology. (5):765.

7. Yao S, Wang S, Zhu Y, Luo L, Chen L. PD-1 on dendritic cells impedes innate immunity against bacterial infection. Blood. 2009;113(23):5811–8.

8. Said EA, Dupuy FP, Trautmann L, Zhang Y, Sekaly RP. Programmed death-1-induced interleukin-10 production by monocytes impairs CD4(+) T cell activation during HIV infection. Nature Medicine. 2010;16(4):452–9.

9. Dong, Haidong, Strome Scott E, Salomao Diva R, et al. Tumor-associated B7-H1 promotes T-cell apoptosis: A potential mechanism of immune evasion. Nature Medicine. 2002.

10. Mojgan, Ahmadzadeh, Laura A, Johnson, Bianca Heemskerk, et al. Tumor antigen-specific CD8 T cells infiltrating the tumor express high levels of PD-1 and are functionally impaired. Blood. 2009.

11. Latchman Y, Wood CR, Chernova T, Chaudhary D, Borde M, Chernova I, et al. PD-L2 is a second ligand for PD-1 and inhibits T cell activation. Nature Immunology. 2001;2(3):261–8.

12. Kim, J. M, Chen, D. S. Immune escape to PD-L1/PD-1 blockade: seven steps to success (or failure). Annals of Oncology. 2016.

13. Han Y, Liu D, Li L. PD-1/PD-L1 pathway: current researches in cancer. American Journal of Cancer Research. 2020;10(3):727–42.

14. Almagro JC, Daniels-Wells TR, Perez-Tapia SM, Penichet ML. Progress and Challenges in the Design and Clinical Development of Antibodies for Cancer Therapy. Frontiers in Immunology. 2017;8:1751.

15. Chames P, Regenmortel MV, Weiss E, Baty D. Therapeutic antibodies: successes, limitations and hopes for the future. British Journal of Pharmacology. 2009;157(2).

16. Leipold D, Prabhu S. Pharmacokinetic and Pharmacodynamic Considerations in the Design of Therapeutic Antibodies. Clin Transl Sci. 2019;12(2):130–9. doi: 10.1111/cts.12597. PubMed PMID: 30414357; PubMed Central PMCID: PMCPMC6440574.

17. Chen X, Song X, Li K, Zhang T. FcγR-Binding Is an Important Functional Attribute for Immune Checkpoint Antibodies in Cancer Immunotherapy. Frontiers in Immunology. 2019;10:-.

18. Deng R, Bumbaca D, Pastuskovas CV, Boswell CA, West D, Cowan KJ, et al. Preclinical pharmacokinetics, pharmacodynamics, tissue distribution, and tumor penetration of anti-PD-L1 monoclonal antibody, an immune checkpoint inhibitor. MAbs. 2016;8(3):593–603. doi: 10.1080/19420862.2015.1136043. PubMed PMID: 26918260; PubMed Central PMCID: PMCPMC4966836.

19. Kurino T, Matsuda R, Terui A, Suzuki H, Kokubo T, Uehara T, et al. Poor outcome with anti-programmed death-ligand 1 (PD-L1) antibody due to poor pharmacokinetic properties in PD-1/PD-L1 blockade-sensitive mouse models. Journal for Immunotherapy of Cancer. 2020;8(1).

20. Gridelli C, Ardizzoni A, Barberis M, Cappuzzo F, Casaluce F, Danesi R, et al. Predictive biomarkers of immunotherapy for non-small cell lung cancer: results from an Experts Panel Meeting of the Italian Association of Thoracic Oncology. Translational Lung Cancer Research. 2017;6(3):373–86.

21. Pierre, Bruhns, Bruno, Iannascoli, Patrick, England, et al. Specificity and affinity of human Fcgamma receptors and their polymorphic variants for human IgG subclasses. Blood. 2009.

22. Gillis C, Gouel-Cheron A, Jonsson F, Bruhns P. Contribution of Human FcgammaRs to Disease with Evidence from Human Polymorphisms and Transgenic Animal Studies. Front Immunol. 2014;5:254. doi: 10.3389/fimmu.2014.00254. PubMed PMID: 24910634; PubMed Central PMCID: PMCPMC4038777.

23. Zhang T, Song X, Xu L, Ma J, Zhang Y, Gong W, et al. The binding of an anti-PD-1 antibody to FcγRΙ has a profound impact on its biological functions. Cancer Immunology, Immunotherapy. 2018.

24. Labrijn, Aran F, Aalberse, Rob C, Bleeker, Wim K, et al. Therapeutic IgG4 antibodies engage in Fab-arm exchange with endogenous human IgG4 in vivo.

25. Swisher Jennifer FA, Feldman Gerald M. The many faces of FcRI: implications for therapeutic antibody function.

26. Nimmerjahn F, Ravetch JV. Fcγ receptors as regulators of immune responses. Nature Publishing Group. 2008;(1).

27. Brandsma AM, Schwartz SL, Wester MJ, Valley CC, Blezer GLA, Vidarsson G, et al. Mechanisms of inside-out signaling of the high-affinity IgG receptor FcgammaRI. Sci Signal. 2018;11(540). doi: 10.1126/scisignal.aaq0891. PubMed PMID: 30042128; PubMed Central PMCID: PMCPMC7521114.

28. Ioan-Facsinay A, Kimpe S, Hellwig S, Lent P, Verbeek JS. FcgammaRI (CD64) contributes substantially to severity of arthritis, hypersensitivity responses, and protection from bacterial infection. Immunity. 2002;16(3):391–402.

29. Shchelokov D, Demin O, editors. Abstract 2233: Prediction and comparison of PD-1 receptor occupancy in the tumor after treatment with immune checkpoint inhibitors. Proceedings: AACR Annual Meeting 2020; April 27-28, 2020 and June 22-24, 2020; Philadelphia, PA; 2020.

30. Wang J, Fei K, Jing H, Wu Z, Wu W, Zhou S, et al. Durable blockade of PD-1 signaling links preclinical efficacy of sintilimab to its clinical benefit. MAbs. 2019;11(8):1443–51. doi: 10.1080/19420862.2019.1654303. PubMed PMID: 31402780; PubMed Central PMCID: PMCPMC6816392.

31. Fu J, Wang F, Dong LH, Xing MJ, Song HF. Receptor occupancy measurement of anti-PD-1 antibody drugs in support of clinical trials. Bioanalysis. 2019;11(72).

32. Brahmer JR, Drake CG, Wollner I, Powderly JD, Picus J, Sharfman WH, et al. Phase I study of single-agent anti-programmed death-1 (MDX-1106) in refractory solid tumors: safety, clinical activity, pharmacodynamics, and immunologic correlates. J Clin Oncol. 2010;28(19):3167–75. doi: 10.1200/JCO.2009.26.7609. PubMed PMID: 20516446; PubMed Central PMCID: PMCPMC4834717.

33. Lou B, Wei H, Yang F, Wang S, Yang B, Zheng Y, et al. Preclinical Characterization of GLS-010 (Zimberelimab), a Novel Fully Human Anti-PD-1 Therapeutic Monoclonal Antibody for Cancer. Front Oncol. 2021;11:736955. doi: 10.3389/fonc.2021.736955. PubMed PMID: 34604074; PubMed Central PMCID: PMCPMC8479189.

34. Kumar S, Ghosh S, Sharma G, Wang Z, Kehry M, Marino M, et al. Preclinical characterization of dostarlimab, a therapeutic anti-PD-1 antibody with potent activity to enhance immune function in in vitro cellular assays and in vivo animal models. mAbs. 13(1):1954136.

35. Sharma S. Pan-TGFβ inhibition by SAR439459 relieves immunosuppression and improves antitumor efficacy of PD-1 blockade. OncoImmunology. 2020;9(1).

